# Flipping the switch on some of the slowest mutating genomes: Direct measurements of plant mitochondrial and plastid mutation rates in *msh1* mutants

**DOI:** 10.1101/2025.01.08.631957

**Authors:** Amanda K. Broz, Mychaela M. Hodous, Yi Zou, Patricia C. Vail, Zhiqiang Wu, Daniel B. Sloan

## Abstract

Plant mitochondrial and plastid genomes have exceptionally slow rates of sequence evolution, and recent work has identified an unusual member of the *MutS* gene family (“plant *MSH1*”) as being instrumental in preventing point mutations in these genomes. However, the effects of disrupting *MSH1*-mediated DNA repair on “germline” mutation rates have not been quantified. Here, we used *Arabidopsis thaliana* mutation accumulation (MA) lines to measure mutation rates in *msh1* mutants and matched wild type (WT) controls. We detected 124 single nucleotide variants (SNVs: 49 mitochondrial and 75 plastid) and 668 small insertions and deletions (indels: 258 mitochondrial and 410 plastid) in *msh1* MA lines. In striking contrast, we did not find any organelle mutations in the WT MA lines, and reanalysis of data from a much larger WT MA experiment also failed to detect any variants. The observed number of SNVs in the *msh1* MA lines corresponds to estimated mutation rates of 6.1×10^-7^ and 3.2 ×10^-6^ per bp per generation in mitochondrial and plastid genomes, respectively. These rates exceed those of species known to have very high mitochondrial mutation rates (e.g., nematodes and fruit flies) by an order of magnitude or more and are on par with estimated rates in humans despite the generation times of *A. thaliana* being nearly 100-fold shorter. Therefore, disruption of a single plant-specific genetic factor in *A. thaliana* is sufficient to erase or even reverse the enormous difference in organelle mutation rates between plants and animals.

## Introduction

Identifying the evolutionary and molecular mechanisms that determine mutation rates remains one of the defining challenges in the field of genetics (Sturtevant 1937; Lynch *et al*. 2016). In addition to the nuclear genome, eukaryotes harbor endosymbiotically derived organelles (mitochondria and plastids) that retain their own genomes. Despite occupying the same cells, these genomic compartments can exhibit highly divergent mutation rates (Brown *et al*. 1979; Wolfe *et al*. 1987; Smith and Keeling 2015). In many eukaryotes (including most animals), the mitochondrial genome has much higher rates of nucleotide substitutions than the nucleus, but the opposite is true in seed plants, which have average nuclear, plastid, and mitochondrial substitution rates that exhibit an approximately 10:3:1 ratio (Drouin *et al*. 2008). Although mitochondrial and plastid mutation rates have not been directly estimated in land plants, phylogenetic analyses indicate that mitochondria experience less than one substitution per site per billion years in many lineages (Mower *et al*. 2007).

One likely explanation for the anomalously low mutation rates in plant organelles is the function of the “plant” *MutS Homolog 1* (*MSH1*) gene, which was horizontally acquired by the green plant lineage (i.e., prior to the divergence of all green algae and land plants) and is absent from most other eukaryotes (Abdelnoor *et al*. 2003; Wu *et al*. 2020). *MSH1* is a member of a much larger gene family with diverse roles in DNA mismatch repair and regulating recombination (Sloan *et al*. 2024). MSH1 is a nuclear-encoded enzyme that is dual-targeted to the mitochondria and plastids and has long been known to maintain the structural stability of plant organelle genomes by suppressing ectopic recombination between small repeated sequences (Martinez-Zapater *et al*. 1992; Abdelnoor *et al*. 2003; Davila *et al*. 2011; Xu *et al*. 2011; Zou *et al*. 2022). The distinctive domain architecture of the MSH1 protein (Abdelnoor *et al*. 2006; Fukui *et al*. 2018; Peñafiel-Ayala *et al*. 2024) has prompted the hypothesis that it is also responsible for maintaining low mutation rates via a novel mismatch repair mechanism (Christensen 2014; Ayala-García *et al*. 2018; Broz *et al*. 2022). This hypothesis is supported by recent findings that disruption of *MSH1* leads to a large increase in the number of *de novo* point mutations (Wu *et al*. 2020; Zou *et al*. 2022; Lencina *et al*. 2022). However, the approaches used in previous studies have precluded direct quantification of the heritable (“germline”) mutation rates in *msh1* mutants.

Mutation accumulation (MA) experiments have proven to be an effective way to generate direct estimates of germline mutation rates in many model systems (Halligan and Keightley 2009; Katju and Bergthorsson 2019; Wang and Obbard 2023). These experiments are conducted by rearing multiple MA lines in the lab. Each generation, lines are propagated by randomly choosing one or a small number of individuals for breeding, thereby minimizing effects of natural selection except in cases of mutations that are completely lethal or sterilizing. Resequencing MA lines after many generations can then identify the *de novo* mutations that have occurred over the course of the experiment. Here, we report the results of an MA experiment, in which we propagated *msh1* mutant lines and matched wild type (WT) control lines in the model angiosperm *Arabidopsis thaliana* to directly quantify mitochondrial and plastid mutation rates.

## Results and Discussion

### Arabidopsis thaliana MA lines

All lines in this study were derived from an *A. thaliana* Col-0 WT plant (maternal parent) that was crossed with a homozygous knock out *msh1* CS3246 mutant line (i.e., *chm1-2*; Abdelnoor *et al*. 2003) to generate heterozygous F1s with a “clean” cytoplasmic background that had never experienced a homozygous *msh1* mutant nuclear genotype (Wu *et al*. 2020). An F1 plant was then self-fertilized to generate F2 families segregating at the *MSH1* locus. F2 plants that were homozygous for WT allele (W1, W2, W3) or the mutant *msh1* allele (M1, M2, M3) were then selected. The progeny from these F2 plants were used to generate the initial material for this study. For each F3 family, seven WT plants (e.g., W1, 1-7) and eight mutant plants (e.g., M1, 1-8) were selected as starting individuals and propagated to the F8 generation (i.e., seven generations in the homozygous WT or *msh1* mutant state) with single-seed descent. Consistent with previous characterizations (Redei 1973; Martinez-Zapater *et al*. 1992; Abdelnoor *et al*. 2003; Virdi *et al*. 2015), individuals from the *msh1* MA lines exhibited a diverse range of mutant phenotypes (Figure 1A). Four *msh1* mutant lines were considered extinct at the F7 generation because F8 seeds would not germinate after multiple plantings, so we analyzed F7 individuals in these cases. Although we began with 21 WT and 24 *msh1* mutant lines, we only included sequence data from 20 WT and 22 *msh1* lines. One WT line was excluded because the sequencing library produced very low yield. Two *msh1* lines were excluded because it was discovered that they had been contaminated by WT seed and were not actually *msh1* mutants.

**Figure 1.**
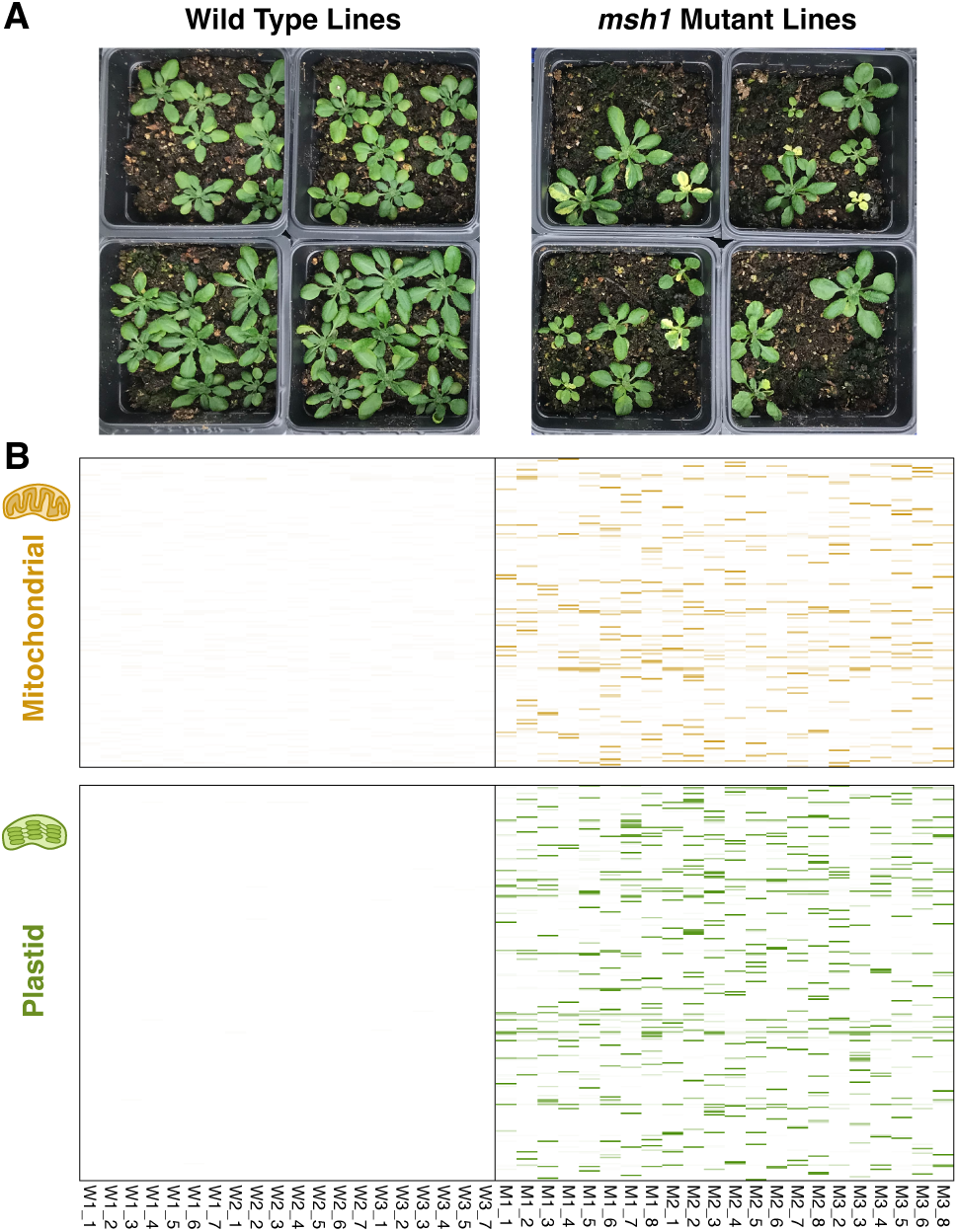
Overview comparison between *A. thaliana* WT (left) and *msh1* mutant (right) MA lines. (A) Examples of phenotypic variation arising in the *msh1* MA lines. F9 progeny from sequenced F8 individuals are pictured. Nine seeds were sown per pot, but not all seeds germinated. (B) Mitochondrial (top) and plastid (bottom) variants detected by MA line sequencing. Each column represents an MA line. Each row represents a nucleotide position in the genome. Only positions with variants are shown, so spacing on the vertical axis is not a measure of distance. The color intensity of horizontal lines in each of cell of the heatmaps represents variant frequency, which was normalized for background sequencing error rates by subtracting the mean variant frequency averaged across all WT MA lines. As indicated by the almost completely white panels, we did not detect any variants that passed filtering criteria in the WT MA lines. We detected 892 SNVs and small indels in the *msh1* mutant MA lines.

### Mitochondrial and plastid germline mutation rates are extremely high in msh1 MA lines

After filtering to exclude variant calling artefacts related to sequencing errors, repeat-mediated recombination, structural rearrangements, and nuclear copies of mitochondrial DNA (numts), we identified a total of 49 mitochondrial single nucleotide variants (SNVs) and 75 plastid SNVs in the 22 *msh1* MA lines. We also observed one mitochondrial multinucleotide variant (MNV) and one plastid MNV, which consisted of either two substitutions at adjacent positions or replacement of a single base pair by a dinucleotide (Dataset S1). We did not identify any organellar SNVs or MNVs in the 20 WT MA lines (Figure 1B). Sanger sequencing of a sample of ten identified SNVs (five mitochondrial and five plastid) confirmed the accuracy of the variant calls and that they were germline mutations transmitted to the F9 generation. One mitochondrial SNV (AT→GC at position 91,017) was shared by two *msh1* MA lines derived from the same F2 parent, suggesting that it may have arisen a single time in that individual and been retained by both of the lines. Every other SNV and MNV was unique to a single MA line. Therefore, the vast majority of the observed variants appear to have arisen independently during MA line propagation.

We found that 7 of the 49 mitochondrial SNVs (14%) and 45 of the 75 plastid SNVs (60%) were in protein-coding, rRNA, or tRNA gene sequences (Table 1). These frequencies closely mirror the percentage of genic sequence in the *A. thaliana* mitochondrial and plastid genomes (10% and 59%, respectively), suggesting that the mutation rates in these functionally important regions were similar to those in intronic/intergenic sequence and that MA line propagation was effective at minimizing selection. Among the variants in protein-coding sequence (CDS), 3 of the 6 mitochondrial SNVs (50%) and 13 of the 35 plastid SNVs (37%) were synonymous changes (i.e. they do not alter the amino acid sequence of the encoded protein). The observed counts of synonymous vs. nonsynonymous variants do not differ significantly from expectations generated by random permutations (*P* = 0.35), again suggesting that selection on functional effects of mutations was very limited in the MA lines.

**Table 1.** Counts of mitochondrial and plastid SNVs observed in *msh1* MA lines by genomic location.

| SNV Location | Mitochondrial | Plastid |
| --- | --- | --- |
| Genic (Total) | 14 | 53 |
| *CDS (synonymous)* | 6 (3) | 35 (13) |
| *rRNA* | 1 | 7 |
| *tRNA* | 0 | 3 |
| *Intronic* | 7 | 8 |
| Intergenic | 35 | 22 |
| **Total** | **49** | **75** |

We detected 50 of the 75 plastid SNVs (67%) at a frequency of 98% or higher in their respective MA line. Allowing for the effects of sequencing and read mapping errors, it is likely that these variants are fixed (homoplasmic) in the sequenced samples. The remaining 25 plastid SNVs are more likely heteroplasmic variants, with measured frequencies ranging from 97% down to the 20% cutoff that was applied during variant detection. Distinguishing between homoplasmic and heteroplasmic SNVs is more challenging for the mitochondrial genome because our sequencing data were generated from total-cellular DNA samples and the *A. thaliana* Col-0 nuclear genome contains a large numt that is nearly identical in sequence to the actual mitochondrial genome (Stupar *et al*. 2001; Fields *et al*. 2022). As such, even if a mitochondrial variant has reached homoplasmy, samples will still produce numt-derived reads that match the reference allele. Therefore, we applied a correction to allele frequency estimates based on the percentage of nuclear sequence in the dataset and the number of copies of the corresponding region in the numt. Given the very approximate nature of this correction, we consider mitochondrial SNVs with an estimated frequency >90% to be strong candidates for homoplasmic variants. Only 14 of the 49 (29%) of the mitochondrial SNVs reached this threshold. The apparently lower proportion of homoplasmic SNVs in the mitochondria compared to the plastids is consistent with previous observations in *A. thaliana msh1* mutants that the plastid genome exhibits stronger effects of transmission bottlenecks and, thus, more rapid heteroplasmic sorting (Broz *et al*. 2022, 2024).

The observed numbers of SNVs correspond to estimated nucleotide substitution rates in *msh1* mutants of 6.1×10^-7^ and 3.2 ×10^-6^ in the mitochondrial and plastid genomes, respectively (here and throughout, mutation rates are expressed as per bp per generation). These values represent remarkably high rates of nucleotide substitution, not only given the low mutations rates that are characteristic of plant organelle genomes but also in comparison to MA line estimates in animal systems with high mitochondrial mutation rates. For example, the mitochondrial nucleotide substitution rates estimated from MA experiments with multiple nematode species and the fruit fly *Drosophila melanogaster* are ∼10-fold and ∼50-fold lower than the mitochondrial and plastid rates we observed in *A. thaliana msh1* lines, respectively (Table 2). Indeed, the observed rates in these *msh1* lines are similar to the germline mitochondrial substitution rates estimated from human pedigrees (Connell *et al*. 2021; Árnadóttir *et al*. 2024), despite the ∼100-fold shorter generation time in *A. thaliana*. To our knowledge, the only other direct estimate of a germline mitochondrial substitution rate that approaches these levels is from the water flea *Daphnia magna* (Ho *et al*. 2020), which intriguingly has an estimated rate that is almost 30-fold higher than that of its congener *D. pulex* (Xu *et al*. 2012).

**Table 2.** Comparison of mitochondrial and plastid mutation rate estimates from published MA and pedigree-based studies. Rates are expressed per base-pair per generation.

| Species | Group | Genome | SNV Rate | Reference |
| --- | --- | --- | --- | --- |
| *Arabidopsis thaliana msh1* | Viridiplantae (land plants) | Mitochondrial | $6.1 \times 10^{-7}$ | This study |
| *Arabidopsis thaliana msh1* | Viridiplantae (land plants) | Plastid | $3.2 \times 10^{-6}$ | This study |
| *Arabidopsis thaliana* WT | Viridiplantae (land plants) | Mitochondrial | Undetected | This study; Weng *et al.* 2019 |
| *Arabidopsis thaliana* WT | Viridiplantae (land plants) | Plastid | Undetected | This study; Weng *et al.* 2019 |
| *Chlamydomonas incerta* | Viridiplantae (chlorophytes) | Mitochondrial | Undetected | López-Cortegano *et al.* 2021 |
| *Chlamydomonas incerta* | Viridiplantae (chlorophytes) | Plastid | $5.4 \times 10^{-10}$ | López-Cortegano *et al.* 2021 |
| *Chlamydomonas reinhardtii* | Viridiplantae (chlorophytes) | Mitochondrial | Undetected | Ness *et al.* 2016 |
| *Chlamydomonas reinhardtii* | Viridiplantae (chlorophytes) | Plastid | $7.7 \times 10^{-10}$ | Ness *et al.* 2016 |
| Mamiellales (4 species) | Viridiplantae (chlorophytes) | Mitochondrial | Undetected | Krasovec *et al.* 2017 |
| Mamiellales (4 species) | Viridiplantae (chlorophytes) | Plastid | Undetected | Krasovec *et al.* 2017 |
| *Phaeodactylum tricornutum* | Stramenopiles (diatoms) | Mitochondrial | $1.1 \times 10^{-9}$ | Krasovec *et al.* 2019 |
| *Phaeodactylum tricornutum* | Stramenopiles (diatoms) | Plastid | Undetected | Krasovec *et al.* 2019 |
| *Paramecium tetraurelia* | Alveolates | Mitochondrial | $7.0 \times 10^{-8}$ | Sung *et al.* 2012 |
| *Dictyostelium discoideum* | Amoebozoa | Mitochondrial | $6.8 \times 10^{-9}$ | Saxer *et al.* 2012 |
| *Saccharomyces cerevisiae* | Fungi | Mitochondrial | $1.2 \times 10^{-8}$ | Lynch *et al.* 2008 |
| *Saccharomyces cerevisiae* | Fungi | Mitochondrial | $2.2 \times 10^{-9}$ | Liu and Zhang 2019 |
| *Caenorhabditis briggsae* | Metazoans (nematodes) | Mitochondrial | $9.1 \times 10^{-8}$ | Howe *et al.* 2010 |
| *Caenorhabditis elegans* | Metazoans (nematodes) | Mitochondrial | $9.7 \times 10^{-8}$ | Denver *et al.* 2000 |
| *Caenorhabditis elegans* | Metazoans (nematodes) | Mitochondrial | $4.3 \times 10^{-8}$ | Konrad *et al.* 2017 |
| *Pristionchus pacificus* | Metazoans (nematodes) | Mitochondrial | $4.5 \times 10^{-8}$ | Molnar *et al.* 2011 |
| *Daphnia magna* | Metazoans (arthropods) | Mitochondrial | $8.7 \times 10^{-7}$ | Ho *et al.* 2020 |
| *Daphnia pulex* | Metazoans (arthropods) | Mitochondrial | $3.2 \times 10^{-8}$ | Xu *et al.* 2012 |
| *Drosophila melanogaster* | Metazoans (arthropods) | Mitochondrial | $6.2 \times 10^{-8}$ | Haag-Liautard *et al.* 2008 |
| *Homo sapiens* | Metazoans (vertebrates) | Mitochondrial | $2.9 \times 10^{-6}$ | Árnadóttir *et al.* 2024 |
| *Homo sapiens* | Metazoans (vertebrates) | Mitochondrial | $1.6 \times 10^{-6}$ | Connell *et al.* 2021 |

Because we did not detect any mitochondrial or plastid SNVs in the *A. thaliana* WT MA lines, we cannot calculate WT mutation rates to compare to the *msh1* lines. To further investigate the occurrence of organelle variants in WT backgrounds, we reanalyzed published data from a much larger *A. thaliana* MA experiment (Weng *et al*. 2019), which resequenced 107 lines at the F25 generation. The combination of more lines and more generations means that this study had a ∼17-fold larger WT sample than our own. Nevertheless, we still did not detect any mitochondrial or plastid SNVs in this dataset, consistent with the lack of reported organelle variants in the original study (Weng *et al*. 2019). To assess a potential upper bound on substitution rates in WT lines, we can consider that if we had observed a single (homoplasmic) SNV in these studies, it would have corresponded to mutation rate estimates of ∼1×10^-9^ in the mitochondrial genome or ∼3×10^-9^ in the plastid genome. More conservatively, we can calculate the lowest mutation rate that would still have yielded a 95% probability (when modeled as a Poisson process) of detecting at least one SNV given the sample size of these studies, which would correspond to rates of ∼3×10^-9^ in the mitochondrial genome and ∼8×10^-9^ in the plastid genome. Given these estimated upper bounds for WT mutation rates, it is not surprising that neither study detected any organelle mutations. Previous phylogenetic analysis of synonymous substitutions has estimated a mitochondrial mutation rate of ∼2×10^-10^ (assuming a generation time of ∼4 months) for the *Arabidopsis* lineage (Mower *et al*. 2007). Based on typical ratios of plastid to mitochondrial substitution rates in angiosperms (Drouin *et al*. 2008), this would correspond to a plastid rate of ∼1×10^-9^. Both these values fall below the measurable range in this study. To our knowledge, the only other direct estimates of mutation rates in photosynthetic eukaryotes have been performed with unicellular species that have much shorter generation times than multicellular plants (Ness *et al*. 2016; Krasovec *et al*. 2017, 2019; López-Cortegano *et al*. 2021). Most of these studies did not detect any mitochondrial or plastid SNVs, and those that did estimated mutation rates of ∼1×10^-9^ or lower (Table 2).

Our earlier study using high-fidelity DNA sequencing to detect low-frequency variants in pooled samples of vegetative tissues estimated that SNV frequencies were ∼10-fold and ∼200-fold higher in *msh1* mutants compared to WT for mitochondrial and plastid genomes, respectively (Wu *et al*. 2020). Despite the large magnitude of these differences, the results from the present study indicate that comparing frequencies of somatic variants may greatly underestimate the proportional effects of disrupting *MSH1* on germline mutation rates. For example, the estimated SNV rate in *msh1* MA lines is ∼200-fold higher than the calculated WT upper bound in the mitochondrial genome and ∼400-fold higher in the plastid genome. Because those ratios are based on WT upper bounds, they represent a minimum estimate of *msh1* mutant effects. Using the phylogenetic-based estimates described above as our WT rates suggests that disrupting *MSH1* increases both the mitochondrial and plastid substitution rate by ∼3000-fold. However, that estimate should be interpreted with caution because phylogenetic analyses have often been found to underestimate mutation rates relative to direct measurements (Parsons *et al*. 1997; Denver *et al*. 2000).

*The mutation spectrum in msh1 MA lines is GC-biased in the plastid genome and GC-neutral in the mitochondrial genome* The observed substitutions in *A. thaliana msh1* mutant lines exhibited a highly biased mutation spectrum (Figure 2). Strikingly, 62 of the 75 (83%) plastid SNVs were AT→GC transitions. This substitution type was also the most common in the mitochondrial genome (23 of 49 SNVs; 47%). The abundance of AT→GC transitions in these lines is consistent with the disproportionate increase in this mutation class that was previously observed among low-frequency somatic variants in *msh1* mutants (Wu *et al*. 2020). In both organelles, GC→AT transitions were the second most common SNV: 6 of 75 (8%) in the plastid genome and 21 of 49 (43%) in the mitochondrial genome. Therefore, transitions represented ∼90% of the observed SNVs in both genomes (Figure 2), but there was a large difference in GC bias between the plastids and the mitochondria. Substitutions were highly GC-biased in the plastid genome, whereas they were relatively GC-neutral in the mitochondrial genome. Even a GC-neutral mutation spectrum is fairly unusual because of the predominant AT bias that has been documented across the tree of life (Hershberg and Petrov 2010; Hildebrand *et al*. 2010). The observed mitochondrial mutation spectrum in *msh1* MA lines is largely congruent with the relatively GC-neutral nucleotide composition of the *A. thaliana* mitochondrial genome (45% GC). In contrast, the GC-biased mutation spectrum in plastids is strikingly opposite the AT-biased composition of the plastid genome (35% GC). Therefore, the loss of *MSH1* function may alter not just the rate but also the spectrum of germline mutations.

**Figure 2.**
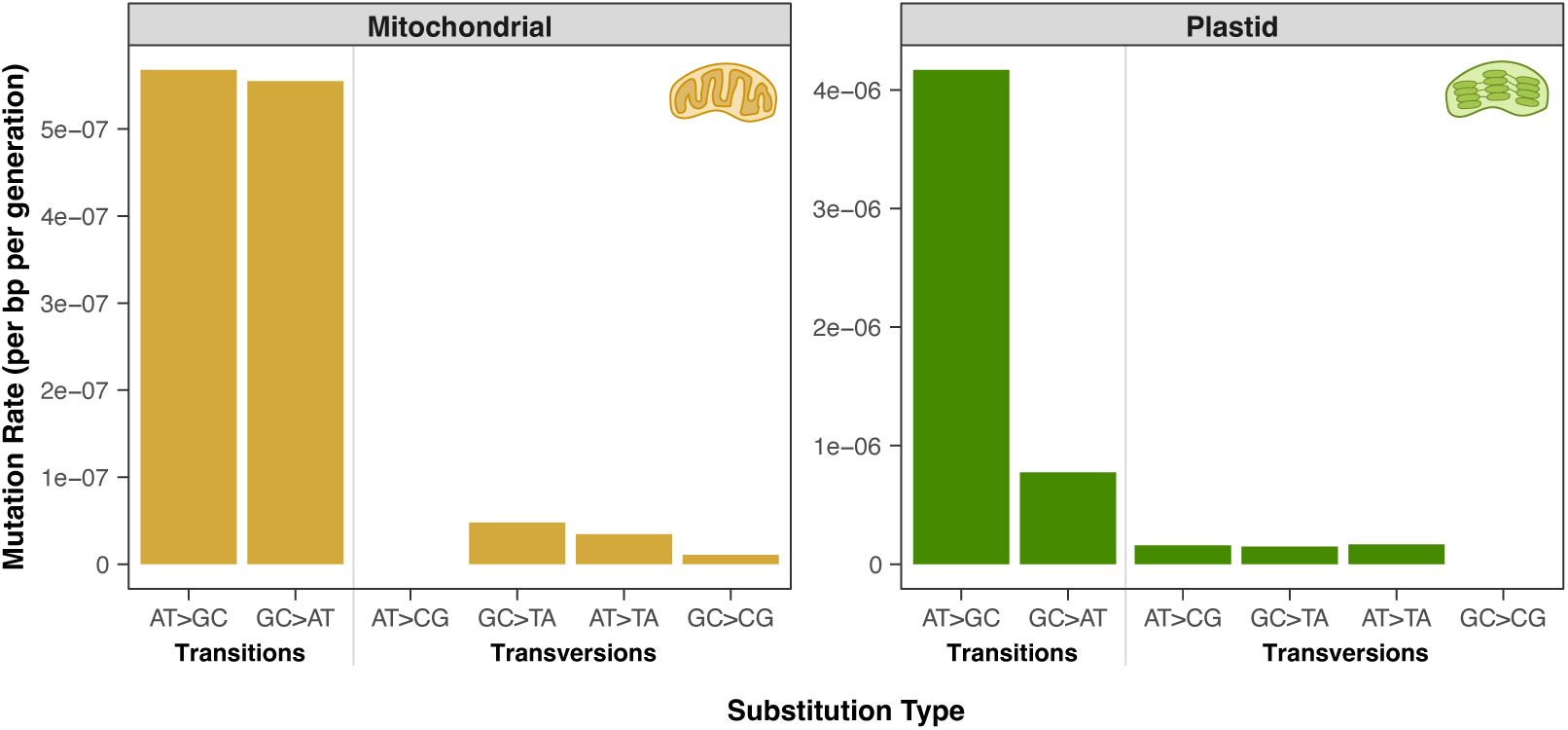
Mitochondrial (left) and plastid (right) mutation spectra in *A. thaliana msh1* MA lines. For mutation rate calculations, SNVs were weighted by heteroplasmic frequency and normalized to the corresponding number of GC or AT base-pairs to account for biased nucleotide compositions in the respective genomes. Note that the y-axis scales differ between the two panels.

We posit three possible explanations (none of which are mutually exclusive) for why the observed mutation spectra in *msh1* MA lines are GC-biased or at least lack the strong AT bias that is often found in other organisms/genomes. First, the extent to which repair pathways are dependent on MSH1 may alter mutation spectra. For example, even if MSH1 targets DNA damage that can cause AT-biased mutations, plant organelles may utilize redundant pathways to repair some of the most common types of damage such as deaminated cytosines (Boesch *et al*. 2009; Córdoba-Cañero *et al*. 2010) and 8-oxoG modifications formed through oxidation (Córdoba-Cañero *et al*. 2014). In contrast, if MSH1 functions in the primary or sole pathway responsible for repair of damage that leads to GC-biased mutations, *msh1* mutants would be expected to show a disproportionate increase in GC-biased mutations. In animal mitochondrial genomes, oxidative damage to adenosines has been hypothesized to cause mutation signatures involving AT→GC transitions (Mikhailova *et al*. 2022). Such damage could potentially be a target of MSH1 in plant organelles. Second, the abundance of AT→GC transitions in the *msh1* mutants may reflect the nucleotide misincorporation kinetics of the DNA polymerases that function in plant organelles. Steady-state kinetic analyses conducted *in vitro* have suggested that these polymerases are prone to misincorporate guanosines opposite thymines in the template strand (Ayala-García *et al*. 2018). If MSH1 normally plays a role in recognition and repair of the resulting mismatches, loss of MSH1 function would again be expected to lead to a disproportionate increase in AT→GC transitions. Finally, it is possible that observed spectra do not solely reflect mutational input but are also biased by selection or a selection-like process. The premise of MA experiments is that the random bottleneck each generation reduces effective population size to an extent that effects of selection are essentially eliminated. However, this experimental bottlenecking is applied at the organismal level and does not preclude selection acting on the large population of organelle genome copies that exists *within* an individual (Taylor *et al*. 2002; Schaack *et al*. 2020). Biased gene conversion can also mimic selection in its effects on allele frequencies (Duret and Galtier 2009). We previously observed that heteroplasmic sorting shows a bias towards GC alleles in *A. thaliana* (Broz *et al*. 2022). Such a form of selection at an intracellular level could bias estimates of both the mutation rate and spectrum relative to true mutational input.

### Homopolymers are hotspots for indel mutations in msh1 MA lines

The effects of disrupting *MSH1* on organelle mutation rates were even more extreme for small indel variants than for SNVs. In the 22 *A. thaliana msh1* mutant MA lines, we detected a total of 258 small indels in the mitochondrial genome and 410 in the plastid genome (Dataset S2). In contrast, we did not detect any small indels in the 20 WT MA lines. The observed small indels were almost exclusively found at homopolymers, with 655 of the 668 small indels (98%) representing expansions or contractions of single-nucleotide repeats of 5-bp or longer. Of the remaining 13 indels, 10 were expansions or contractions of tandem dinucleotide repeats. Therefore, replication of simple repetitive sequences appears to be extremely error-prone in the absence of MSH1 function. The extreme mutability of these loci likely explains why there were numerous cases where mutations were detected at the same homopolymer or dinucleotide repeat in multiple MA lines (Dataset S2). Although some of these may represent mutations that arose in an F2 progenitor and were inherited by multiple lines, we infer that the vast majority represent “multiple hits” that occurred independently in the different lines. This conclusion is supported by the following evidence: (1) it is similarly common for lines derived from different F2 individuals share the same small indel, (2) there are many cases where the same homopolymer exhibits mutations in multiple lines that differ in indel length, and (3) sharing of alleles among lines was extremely rare for SNVs (see above).

The observed counts of small indels correspond to exceptionally high mutation rates of 4.8×10^-6^ and 2.1×10^-5^ in mitochondrial and plastid genomes, respectively. Because of the high rates of sequencing errors and other challenges to accurately estimating indel allele frequency at homopolymers, we did not attempt to distinguish between heteroplasmic and homoplasmic indel variants. Therefore, the above rate estimates may be inflated because we did not weight variants by their allele frequencies. On balance, however, these values are more likely to be underestimates given our inability to capture multiple independent mutations at the same site in a single line or variants with frequencies that do not rise above the high noise threshold due to sequencing error rates at homopolymers.

The higher observed rate of small indel mutations in plastids than in mitochondria may largely reflect the greater abundance of homopolymers in the plastid genome, especially the large number of long A/T homopolymers. The homopolymer composition of the respective genomes may also influence the balance of insertion vs. deletion mutations. Both genomes exhibit a bias towards deletions at A/T homopolymers and a bias towards insertions at G/C homopolymers (Figure 3A). The AT-rich plastid genome has 12.3-fold more A/T homopolymers than G/C homopolymers (Figure 3B). Accordingly, its overall indel spectrum is deletion-biased (140 insertions and 270 deletions). In the mitochondrial genome, there are only 2.8-fold more A/T homopolymers than G/C homopolymers, and the overall indel spectrum is slightly insertion-biased (144 insertions and 114 deletions). These mutation patterns are an illustration of the reciprocal causal relationships that can exist between genome nucleotide composition and mutation rates/biases and potentially lead to feedback cycles.

**Figure 3.**
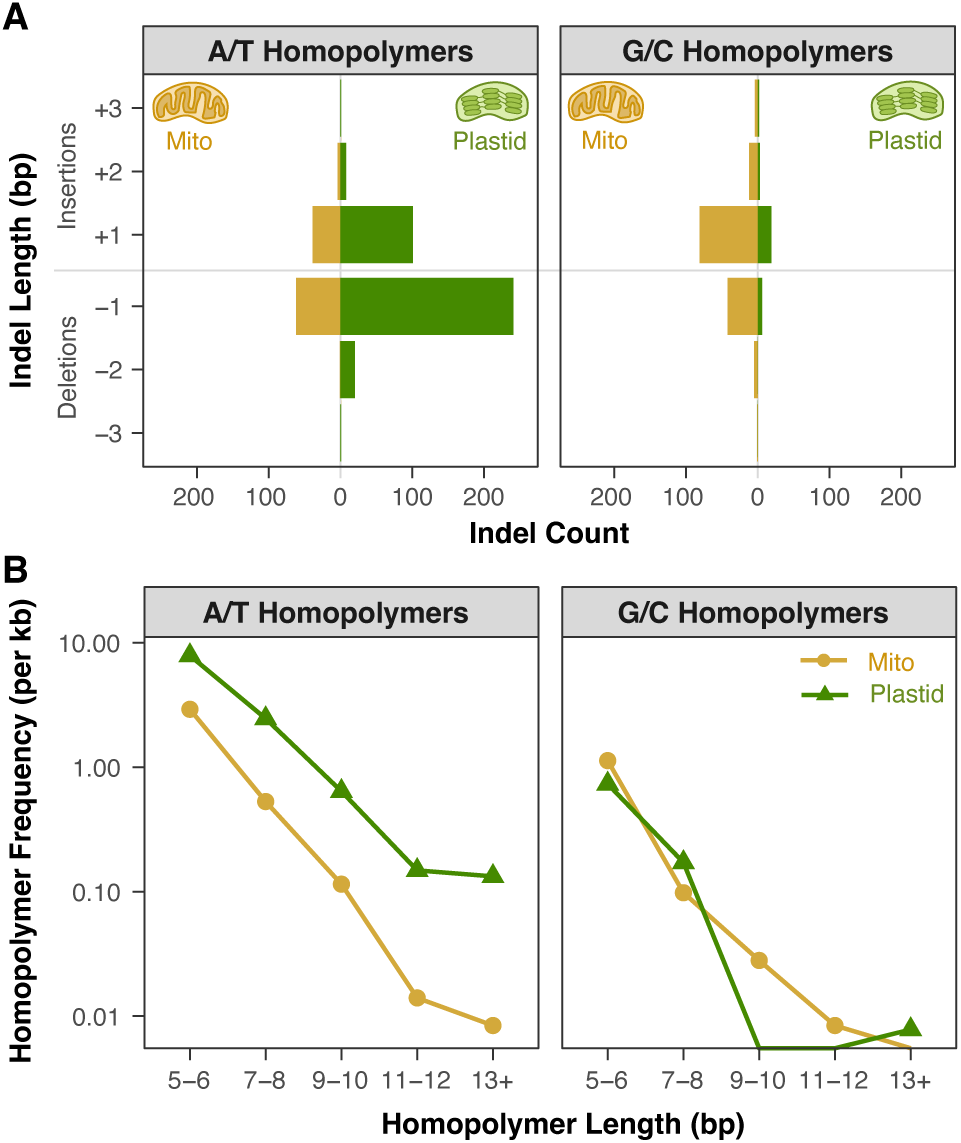
The relationship between homopolymer types and indel bias in *A. thaliana msh1* MA lines. (A) The bars represent the counts of short indels observed in A/T homopolymers (left panel) and G/C homopolymers (right panel). Within each panel, mitochondrial indels are shown to the left of the center line, and plastid indels are shown to the right. Both genomes exhibit a deletion bias in A/T homopolymers and an insertion bias in G/C homopolymers. Indels >3 bp in length were extremely rare (Dataset S3) and not shown in this plot. (B) Frequency of A/T homopolymers (left panel) and G/C homopolymers (right panel) in the *A. thaliana* mitochondrial (gold circles) and plastid (green triangles) genomes. The fact that the plastid genome is dominated by A/T homopolymers may explain why its overall indel spectrum is deletion-biased given that A/T homopolymers appear to be more deletion-prone than G/C homopolymers in both organelles of *msh1* MA lines.

A previous study that identified the spontaneous plastid mutations responsible for observed phenotypes in collections of the evening primrose *Oenothera* also found a predominance of small indels (Massouh *et al*. 2016). However, those variants did not exhibit the same overwhelming effect of homopolymer runs that we observed in *A. thaliana msh1* MA lines, suggesting that the loss of MSH1 function may preferentially exacerbate error rates at homopolymers.

### WT and msh1 MA lines show little diGerence in nuclear mutation rates

Given the known targeting of MSH1 to the mitochondria and chloroplasts, we anticipated that the mutagenic effects of disrupting its function would be limited to the organelle genomes. To test this prediction, we analyzed our sequence data to identify *de novo* nuclear mutations, filtering for SNVs that were found in only a single MA line and determined to be homozygous. Interestingly, the mean number of nuclear SNVs per line was significantly higher for *msh1* mutants than for WT lines (7.0 vs 3.4; *P* = 0.006; Dataset S3). Although this difference is tiny in comparison to the increases in mutation rate observed in the organelle genomes, it could indicate that there is a mutagenic effect in the nuclear genome resulting from the loss of MSH1 function. For example, it has been shown that *msh1* mutants exhibit altered cytosine methylation patterns in the nuclear genome perhaps due to retrograde signaling from the plastids (Virdi *et al*. 2015; Kundariya *et al*. 2022), which could potentially affect the rate and locations of mutations. In addition, the highly disruptive effects on organelle genomes in *msh1* mutants may alter plant biochemistry, physiology, and growth (Figure 1A) with indirect effects on the nuclear mutation rate. However, the apparent difference between *msh1* mutant and WT lines in nuclear mutation rate should be interpreted with caution, particularly because it appears to be driven by lower rates in just two of the three sets of WT lines. The lines derived from one of the WT F2 plants (W2) show nuclear SNV counts that are comparable to those of the three *msh1* sets (Dataset S3). Given that our analysis filtered for unique nuclear SNVs, it is difficult to explain why there would be an effect of the F2 progenitor, although it is possible that there were sequence variants that distinguished the F2 individuals and affected nuclear mutation rates. Regardless, the effects of *MSH1* on nuclear mutation rates, if any, appear minimal compared to the massive increase in mitochondrial and plastid mutagenesis.

## Conclusions

It has long been perplexing why plants and animals can differ by orders of magnitude in their germline organelle mutation rates despite their similarity in effective population size and other biological features (Lynch 2010). Our study shows that disruption of a single genetic factor in plants (*MSH1*) is sufficient to erase or even reverse this difference. The fact that *MSH1* appears to have been horizontally transferred into the green plant lineage illustrates how acquisition of foreign DNA repair machinery can have a fundamental effect on mutation rate evolution, which has also been observed for other members of the *MutS* gene family with mitochondrial functions (Muthye and Lavrov 2021; Sloan *et al*. 2024). The mechanisms by which MSH1 suppresses substitutions and small indels in plant organelle genomes have not yet been fully determined, but there is some evidence supporting a process involving mismatch recognition followed by introduction of double-stranded DNA breaks and template-based recombinational repair (Christensen 2014; Ayala-García *et al*. 2018; Wu *et al*. 2020; Broz *et al*. 2022; Peñafiel-Ayala *et al*. 2024). Regardless of the mechanism, *MSH1* function appears to be critical for maintaining plant viability over generational timescales. When bottlenecking during MA line propagation limits the efficacy of selection, the accumulation of deleterious mutations was so fast in *msh1* mutant lines that we began to see line “extinctions” in only a handful of generations. Even in natural populations where selection is more efficacious, sustaining organelle function in the face of the extreme rates of nucleotide substitutions, indels, and structural mutations that occur in the absence of MSH1 function would seem untenable.

## Methods

### Plant growth and MA line propagation

The original F2 families used for this experiment were those created in Wu *et al*. (2020). For MA line propagation, seeds were placed in water and vernalized at 4 °C for three days. Seeds were then transferred to 3×3 inch pots filled with PRO-MIX BX potting media and placed on light shelves with a 16-hr day length (light intensity of ∼150 μE/m^2^/s). Nine seeds from each line were sown, and a single randomly chosen individual from each line was allowed to set seed for the next generation. If the randomly chosen individual failed to germinate/survive/reproduce, a backup individual was chosen at random (and so on). This process of single-seed descent was carried out until the F8 generation. Seeds for each subsequent generation were planted within one month of parental seed harvesting.

### DNA extraction and sequencing

A single rosette leaf (or two in the case of very small plants) was harvested from each F8 (or F7) plant prior to bolting and stored at -80 °C until DNA was extracted using a Qiagen DNeasy Plant Kit and quantified with a Qubit HS-dsDNA kit. We performed a pilot round of DNA sequencing on samples from three MA lines (M1_2, M2_1, and W1_1) to assess whether there were likely to be enough variants to robustly measure mutation rates. For this pilot round, Illumina library construction was performed by Novogene, using an NEBNext Ultra II DNA Library Prep Kit (E7645L) and up to 80 ng of input DNA. After the pilot run, libraries for the remaining MA lines were also generated by Novogene but with an ABclonal Rapid Plus DNA Lib Prep Kit for Illumina (RK20208) and 70 ng of input DNA for each sample. In both cases, sequencing was performed on the Novaseq X Plus platform with 2×150bp paired-end reads on a 25B flow cell (partial lane sequencing to generate ≥10 Gb of data per sample).

### Mitochondrial and plastid variant calling

Illumina reads were processed with Cutadapt v4.0 to remove Illumina adapters and low quality end sequence (-q 20), applying a minimum trimmed read length of 50 bp. Trimmed reads were then mapped to the *A. thaliana* Col-0 mitochondrial (Sloan *et al*. 2018) and plastid (Sato *et al*. 1999) genomes using Bowtie v2.2.5 (Langmead and Salzberg 2012). The plastid genome reference was modified to reflect the previous observation that our Col-0 lab stock differs from the published sequence by a 1-bp expansion in the homopolymer at position 28,673 (Wu *et al*. 2020). The resulting alignments were then sorted, converted to bam format, and indexed with Samtools v1.17 (Li *et al*. 2009). Variant counts at each position in the genomes were then compiled with Perbase v 0.9.0 (https://github.com/sstadick/perbase).

Using custom scripts, variants were filtered to only include those with a frequency of ≥20% and coverage depth of ≥50×. We used this relatively high frequency of cutoff of 20% to minimize false positives or misidentify low-frequency vegetative mutations as germline variants. The high cutoff was also important because some low-complexity regions such as long homopolymers cause high sequencing error rates and elevated variant frequencies across all samples, so we further filtered variants to exclude those that did not have at least a three-fold higher frequency than the mean across all WT lines. Plant organelle genomes contain large repeated sequences that are identical in sequence and frequently interconvert via recombination and gene conversion (Arrieta-Montiel and Mackenzie 2011; Zhu *et al*. 2016). To avoid double counting variants in these regions, we removed calls in one copy of the large inverted repeat in the plastid genome and one copy of each of the two pairs of large repeats in the mitochondrial genome.

Disruption of MSH1 function leads to recombinational activity between small, imperfect repeats, resulting in structural rearrangements (Martinez-Zapater *et al*. 1992; Abdelnoor *et al*. 2003; Davila *et al*. 2011; Zou *et al*. 2022). When mapped to a reference sequence, reads from these rearranged genomes can lead to artefactual identification of *de novo* SNVs and small indels (Davila *et al*. 2011; Wu *et al*. 2020). Therefore, we used BLASTN 2.14.1+ (Camacho *et al*. 2009) searches to identify and exclude variants that could be explained by structural rearrangements. Variants were similarly searched against the large numt sequence (Fields *et al*. 2022) to exclude spurious calls related to this insertion in the nuclear genome (Waneka *et al*. 2024). We also corrected estimates of mitochondrial SNV frequencies to account for numt-derived sequencing reads that are identical to the reference allele in the mitochondrial genome. Average nuclear genome coverage was estimated based on the percentage of reads that were not mapped to the organelle genomes by Bowtie 2 and the total size of the nuclear genome. Using this coverage estimate and the number of copies of the corresponding region that are present in the numt, we subtracted the expected number of nuclear-derived reads from both the reference allele count and the total coverage of the mitochondrial locus and then recalculated the variant frequency.

To expand sampling of WT lines, we reanalyzed previously published *A. thaliana* MA line resequencing data (Weng *et al*. 2019) with the same variant calling approach. Although the raw variant calls included 231 SNVs (and no indels) that passed our coverage and frequency thresholds, these were exclusively found in just six of the 107 MA lines (39, 40, 100, 101, 102, and 103), and all of them were present <50% frequency. Moreover, their frequencies were highly correlated across the six sequencing libraries despite the fact that these lines were propagated independently for 25 generations. Therefore, these variants appear to be the result of a biased sequencing error profile in these six libraries, and we concluded that there were no convincing organelle sequence variants in any of these MA lines, consistent with the lack of any identified variants in the original publication (Weng *et al*. 2019).

### Mitochondrial and plastid variant confirmation

To confirm high frequency SNVs and determine whether they are germline variants transmitted to offspring, Sanger sequencing was performed (Genewiz, Azenta Life Sciences) using locus-specific primers (Table S1). Sequenced samples included the F8 individual harboring the SNV, an F9 progeny of this plant (growing conditions as described above in “Plant growth and MA line propagation”), and a WT control line (not harboring the SNV). Sequencing traces were analyzed using Chromas Lite software.

### Mutation Rate Calculations

Germline mutation rates (μ) were calculated as follows:

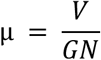

where *V* is the number of identified variants, *G* is the respective genome size, and *N* is the total number of generations summed across the set of MA lines. Genome sizes were reduced by the length of the large repeat copies that were excluded for variant calling purposes (see above), resulting in sizes of 357,025 and 128,214 bp for the mitochondrial and plastid genomes, respectively. There was a total of 150 generations for homozygous *msh1* mutants (7 generations for each of the 18 F8 lines and 6 generations for each of the 4 F7 lines; the F1 generation was excluded from this count because that individual was not homozygous for the *msh1* mutant allele). For SNV mutation rates, each variant was scaled by its frequency. However, due to the high-sequencing error rate and other challenges associated with estimating frequencies at homopolymers, we used unscaled variant counts to calculate indel mutation rates (see above for discussion of associated effects on rate estimates). To calculate an upper bound for the WT mutation rates, we used the fact that the probability of observing zero mutations given a Poisson distribution is e^-λ^, where λ is the expected number of mutations and can be calculated as the product of μ, *G*, and *N*. By setting this probability to 0.05 and solving for μ, the minimum mutation rate that would have yielded a 95% probability of detecting at least one mutation can be calculated as follows:

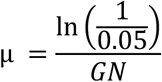

For these upper-bound calculations, *N* was set to 2835, reflecting the sum of 2675 (107 MA lines × 25 generations per line) from Weng *et al*. (2019) and 160 (20 WT MA lines × 8 generations per line) from this study.

### Functional annotation and simulations of nonsynonymous and synonymous variants

Functional characterization of the location of identified variants (genic/intronic/intergenic) and the effect on protein coding sequence (synonymous/nonsynonymous) was conducted with custom scripts. In addition, we simulated random sets of mutations to determine whether the number of synonymous vs. nonsynonymous mutations in our dataset exhibited signatures of selection against deleterious variants. To control for the mutation spectrum, we kept the number (six in the mitochondrial genome and 35 in the plastid genome) and type (AT→GC, GC→AT, GC→TA, etc.) constant but randomized their position within protein-coding sequences. Using 10,000 of these permutations, we generated a distribution of the total count of synonymous substitutions, which we used to perform a one-tailed test for an enrichment of synonymous SNVs in our dataset. We calculated the *P*-value for this test as the frequency of permutations with greater than or equal to the observed number of synonymous SNVs (16) in the dataset.

### Mitochondrial and plastid genome homopolymer analysis

A custom script was used to extract the number, length, and type of homopolymers in the *A. thaliana* mitochondrial and plastid genomes. For consistency with variant calling and mutation rate calculations, reported homopolymer data exclude one copy of each pair of large repeats in these genomes.

### Nuclear variant calling

To identify nuclear SNVs, BWA-MEM v0.7.18-r1243 (Li 2013) was used to map previously trimmed reads (see above) from each of our MA lines to the current *A. thaliana* Col-CC nuclear genome assembly (NCBI GCA_028009825.2) along with the above organelle reference sequences (Reiser *et al*. 2024). GATK v4.6.1.0 MarkDuplicates, HaplotypeCaller, and GenotypeGVCFs were used to identify variants in the resulting alignments, which were filtered with custom scripts to identify homozygous SNVs that had a coverage of >20× and were unique to a single MA line. These variants were further filtered to remove clusters of adjacent SNVs and indels that likely arose from mapping artefacts related to larger structural variants.

## Data and Code Availability

Illumina sequencing reads were deposited to the NCBI Sequencing Read Archive and are available under BioProject PRJNA1201229. Custom scripts used in data analysis and figure generations are available via GitHub (https://github.com/dbsloan/msh1_MA_lines).

## Acknowledgements

We thank Mao-Lun Weng for helpful information about previously sequenced MA lines. This work was supported by funding from the National Institutes of Health (R35GM148134).

---

**Table S1.**
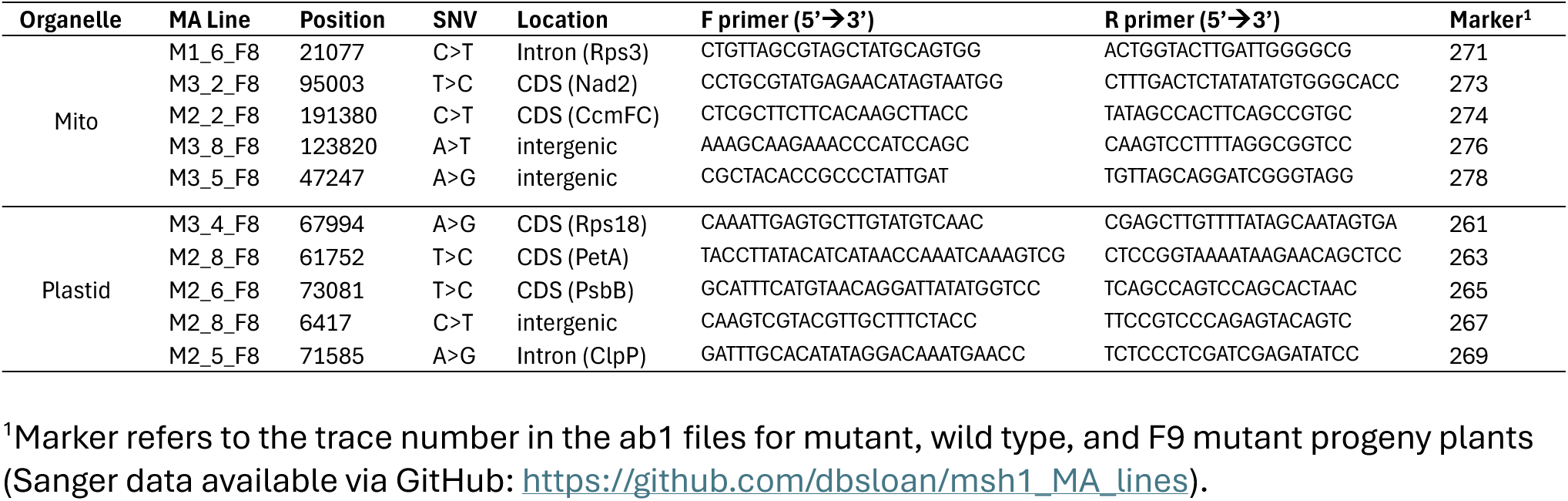
Primers used to validate a sample of mitochondrial and plastid SNVs by PCR amplification and Sanger sequencing. All tested sequences were confirmed as germline variants.

**Dataset S1**. Summary of each organelle SNV.

**Dataset S2**. Summary of each organelle indel.

**Dataset S3**. Summary of each nuclear SNV.

## Notes

### Competing Interest Statement

The authors have declared no competing interest.

## References

Abdelnoor R. V., R. Yule, A. Elo, A. C. Christensen, G. Meyer-Gauen, et al., 2003 Substoichiometric shifting in the plant mitochondrial genome is influenced by a gene homologous to MutS. Proceedings of the National Academy of Sciences 100: 5968–5973.

Abdelnoor R. V., A. C. Christensen, S. Mohammed, B. Munoz-Castillo, H. Moriyama, et al., 2006 Mitochondrial genome dynamics in plants and animals: convergent gene fusions of a MutS homologue. J. Mol. Evol. 63: 165–173.

Árnadóttir E. R., K. H. S. Moore, V. B. Guðmundsdóttir, S. S. Ebenesersdóttir, K. Guity, et al., 2024 The rate and nature of mitochondrial DNA mutations in human pedigrees. Cell 187: 3904–3918.e8.

Arrieta-Montiel M. P., and S. A. Mackenzie, 2011 Plant mitochondrial genomes and recombination, pp. 65–82 in Plant Mitochondria, edited by Kempken F. Springer Verlag, New York.

Ayala-García V. M., N. Baruch-Torres, P. L. García-Medel, and L. G. Brieba, 2018 Plant organellar DNA polymerases paralogs exhibit dissimilar nucleotide incorporation fidelity. FEBS J. 285: 4005–4018.

Boesch P., N. Ibrahim, F. Paulus, A. Cosset, V. Tarasenko, et al., 2009 Plant mitochondria possess a short-patch base excision DNA repair pathway. Nucleic Acids Res. 37: 5690–5700.

Brown W. M., M. George, and A. C. Wilson, 1979 Rapid evolution of animal mitochondrial DNA. Proceedings of the National Academy of Sciences 76: 1967–1971.

Broz A. K., A. Keene, M. Fernandes Gyorfy, M. Hodous, I. G. Johnston, et al., 2022 Rapid sorting of plant mitochondrial and plastid heteroplasmies depends on MSH1 activity. Proceedings of the National Academy of Sciences 119: e2206973119.

Broz A. K., D. B. Sloan, and I. G. Johnston, 2024 Stochastic organelle genome segregation through Arabidopsis development and reproduction. New Phytol. 241: 896–910.

Camacho C., G. Coulouris, V. Avagyan, N. Ma, J. Papadopoulos, et al., 2009 BLAST+: architecture and applications. BMC Bioinformatics 10: 421.

Christensen A. C., 2014 Genes and junk in plant mitochondria-repair mechanisms and selection. Genome Biol. Evol. 6: 1448–1453.

Connell J., M. Benton, R. A. Lea, H. Sutherland, J. Chaseling, et al., 2021 Pedigree derived mutation rate across the entire mitochondrial genome of the Norfolk Island population. Sci. Rep. 12. 10.1038/s41598-022-10530-3

Córdoba-Cañero D., E. Dubois, R. R. Ariza, M. P. Doutriaux, and T. Roldan-Arjona, 2010 Arabidopsis uracil DNA glycosylase (UNG) is required for base excision repair of uracil and increases plant sensitivity to 5-fluorouracil. J. Biol. Chem. 285: 7475–7483.

Córdoba-Cañero D., T. Roldán-Arjona, and R. R. Ariza, 2014 Arabidopsis ZDP DNA 3′-phosphatase and ARP endonuclease function in 8-oxoG repair initiated by FPG and OGG 1 DNA glycosylases. Plant J. 79: 824–834.

Davila J. I., M. P. Arrieta-Montiel, Y. Wamboldt, J. Cao, J. Hagmann, et al., 2011 Double-strand break repair processes drive evolution of the mitochondrial genome in Arabidopsis. BMC Biol. 9: 64.

Denver D. R., K. Morris, M. Lynch, L. L. Vassilieva, and W. K. Thomas, 2000 High direct estimate of the mutation rate in the mitochondrial genome of Caenorhabditis elegans. Science 289: 2342–2344.

Drouin G., H. Daoud, and J. Xia, 2008 Relative rates of synonymous substitutions in the mitochondrial, chloroplast and nuclear genomes of seed plants. Mol. Phylogenet. Evol. 49: 827–831.

Duret L., and N. Galtier, 2009 Biased Gene Conversion and the Evolution of Mammalian Genomic Landscapes. Annu. Rev. Genom. Hum. Genet. 10: 285–311.

Fields P. D., G. Waneka, M. Naish, M. C. Schatz, I. R. Henderson, et al., 2022 Complete Sequence of a 641-kb Insertion of Mitochondrial DNA in the Arabidopsis thaliana Nuclear Genome. Genome Biol. Evol. 14: evac059.

Fukui K., A. Harada, T. Wakamatsu, A. Minobe, K. Ohshita, et al., 2018 The GIY-YIG endonuclease domain of Arabidopsis MutS homolog 1 specifically binds to branched DNA structures. FEBS Lett. 592: 4066–4077.

Haag-Liautard C., N. Coffey, D. Houle, M. Lynch, B. Charlesworth, et al., 2008 Direct estimation of the mitochondrial DNA mutation rate in Drosophila melanogaster. PLoS Biol. 6: 1706–1714.

Halligan D. L., and P. D. Keightley, 2009 Spontaneous mutation accumulation studies in evolutionary genetics. Annu. Rev. Ecol. Evol. Syst. 40: 151–172.

Hershberg R., and D. A. Petrov, 2010 Evidence that mutation is universally biased towards AT in bacteria. PLoS Genet. 6: e1001115.

Hildebrand F., A. Meyer, and A. Eyre-Walker, 2010 Evidence of selection upon genomic GC-content in bacteria. PLoS Genet. 6: e1001107.

Ho E. K. H., F. Macrae, L. C. Latta, P. McIlroy, D. Ebert, et al., 2020 High and highly variable spontaneous mutation rates in Daphnia. Mol. Biol. Evol. In Press.

Howe D. K., C. F. Baer, and D. R. Denver, 2010 High rate of large deletions in Caenorhabditis briggsae mitochondrial genome mutation processes. Genome Biol. Evol. 2: 29–38.

Katju V., and U. Bergthorsson, 2019 Old Trade, New Tricks: Insights into the Spontaneous Mutation Process from the Partnering of Classical Mutation Accumulation Experiments with High-Throughput Genomic Approaches, (K. Makova, Ed.). Genome Biology and Evolution 11: 136–165.

Konrad A., O. Thompson, R. H. Waterston, D. G. Moerman, P. D. Keightley, et al., 2017 Mitochondrial mutation rate, spectrum and heteroplasmy in Caenorhabditis elegans spontaneous mutation accumulation lines of differing population size. Mol. Biol. Evol. 34: 1319–1334.

Krasovec M., A. Eyre-Walker, S. Sanchez-Ferandin, and G. Piganeau, 2017 Spontaneous mutation rate in the smallest photosynthetic eukaryotes. Mol. Biol. Evol. 34: 1770–1779.

Krasovec M., S. Sanchez-Brosseau, and G. Piganeau, 2019 First estimation of the spontaneous mutation rate in diatoms. Genome Biol. Evol. 11: 1829–1837.

Kundariya H., R. Sanchez, X. Yang, A. Hafner, and S. A. Mackenzie, 2022 Methylome decoding of RdDM-mediated reprogramming effects in the Arabidopsis MSH1 system. Genome Biol. 23: 167.

Langmead B., and S. L. Salzberg, 2012 Fast gapped-read alignment with Bowtie 2. Nat. Methods 9: 357– 359.

Lencina F., A. Landau, and A. R. Prina, 2022 The barley chloroplast mutator (cpm) mutant: all roads lead to the Msh1 gene. Int. J. Mol. Sci. 23: 1814.

Li H., B. Handsaker, A. Wysoker, T. Fennell, J. Ruan, et al., 2009 The Sequence Alignment/Map format and SAMtools. Bioinformatics 25: 2078–2079.

Li H., 2013 Aligning sequence reads, clone sequences and assembly contigs with BWA-MEM. arXiv [q-bio.GN].

Liu H., and J. Zhang, 2019 Yeast spontaneous mutation rate and spectrum vary with environment. Curr. Biol. 29: 1584–1591.e3.

López-Cortegano E., R. J. Craig, J. Chebib, T. Samuels, A. D. Morgan, et al., 2021 De Novo mutation rate variation and its determinants in Chlamydomonas. Mol. Biol. Evol. 38: 3709–3723.

Lynch M., W. Sung, K. Morris, N. Coffey, C. R. Landry, et al., 2008 A genome-wide view of the spectrum of spontaneous mutations in yeast. Proc. Natl. Acad. Sci. U. S. A. 105: 9272–9277.

Lynch M., 2010 Evolution of the mutation rate. Trends Genet. 26: 345–352.

Lynch M., M. S. Ackerman, J.-F. Gout, H. Long, W. Sung, et al., 2016 Genetic drift, selection and the evolution of the mutation rate. Nat. Rev. Genet. 17: 704–714.

Martinez-Zapater J. M., P. Gil, J. Capel, and C. R. Somerville, 1992 Mutations at the Arabidopsis CHM locus promote rearrangements of the mitochondrial genome. Plant Cell 4: 889–899.

Massouh A., J. Schubert, L. Yaneva-Roder, E. S. Ulbricht-Jones, A. Zupok, et al., 2016 Spontaneous chloroplast mutants mostly occur by replication slippage and show a biased pattern in the plastome of Oenothera. Plant Cell 28: 911–929.

Mikhailova A. G., A. A. Mikhailova, K. Ushakova, E. O. Tretiakov, D. Iliushchenko, et al., 2022 A mitochondria-specific mutational signature of aging: increased rate of A> G substitutions on the heavy strand. Nucleic acids research 50: 10264–10277.

Molnar R. I., G. Bartelmes, I. Dinkelacker, H. Witte, and R. J. Sommer, 2011 Mutation rates and intraspecific divergence of the mitochondrial genome of Pristionchus pacificus. Mol. Biol. Evol. 28: 2317–2326.

Mower J. P., P. Touzet, J. S. Gummow, L. F. Delph, and J. D. Palmer, 2007 Extensive variation in synonymous substitution rates in mitochondrial genes of seed plants. BMC Evol. Biol. 7: 135.

Muthye V., and D. V. Lavrov, 2021 Multiple losses of MSH1, gain of mtMutS, and other changes in the MutS family of DNA repair proteins in animals. Genome Biol. Evol. 13: evab191.

Ness R. W., S. A. Kraemer, N. Colegrave, and P. D. Keightley, 2016 Direct estimate of the spontaneous mutation rate uncovers the effects of drift and recombination in the Chlamydomonas reinhardtii Plastid genome. Mol. Biol. Evol. 33: 800–808.

Parsons T. J., D. S. Muniec, K. Sullivan, N. Woodyatt, R. Alliston-Greiner, et al., 1997 A high observed substitution rate in the human mitochondrial DNA control region. Nat Genet 15: 363–368.

Peñafiel-Ayala A., A. Peralta-Castro, J. Mora-Garduño, P. García-Medel, A. G. Zambrano-Pereira, et al., 2024 Plant organellar MSH1 is a displacement loop–specific endonuclease. Plant and Cell Physiology 65: 560–575.

Redei G. P., 1973 Extra-chromosomal mutability determined by a nuclear gene locus in Arabidopsis. Mutat. Res. 18: 149–162.

Reiser L., E. Bakker, S. Subramaniam, X. Chen, S. Sawant, et al., 2024 The Arabidopsis Information Resource in 2024. Genetics 227: iyae027.

Sato S., Y. Nakamura, T. Kaneko, E. Asamizu, and S. Tabata, 1999 Complete structure of the chloroplast genome of Arabidopsis thaliana. DNA Res. 6: 283–290.

Saxer G., P. Havlak, S. A. Fox, M. A. Quance, S. Gupta, et al., 2012 Whole genome sequencing of mutation accumulation lines reveals a low mutation rate in the social amoeba Dictyostelium discoideum. PLoS One 7: e46759.

Schaack S., E. K. H. Ho, and F. Macrae, 2020 Disentangling the intertwined roles of mutation, selection and drift in the mitochondrial genome. Phil. Trans. R. Soc. B 375: 20190173.

Sloan D. B., Z. Wu, and J. Sharbrough, 2018 Correction of persistent errors in Arabidopsis reference mitochondrial genomes. Plant Cell 30: 525–527.

Sloan D. B., A. K. Broz, S. A. Kuster, V. Muthye, A. Peñafiel-Ayala, et al., 2024 Expansion of the MutS gene family in plants. Plant Cell In Press. 10.1101/2024.07.17.603841

Smith D. R., and P. J. Keeling, 2015 Mitochondrial and plastid genome architecture: Reoccurring themes, but significant differences at the extremes. Proceedings of the National Academy of Sciences 112: 10177–10184.

Stupar R. M., J. W. Lilly, C. D. Town, Z. Cheng, S. Kaul, et al., 2001 Complex mtDNA constitutes an approximate 620-kb insertion on Arabidopsis thaliana chromosome 2: implication of potential sequencing errors caused by large-unit repeats. Proc. Natl. Acad. Sci. U. S. A. 98: 5099–5103.

Sturtevant A. H., 1937 Essays on Evolution. I. On the Effects of Selection on Mutation Rate. Q. Rev. Biol. 12: 464.

Sung W., A. E. Tucker, T. G. Doak, E. Choi, W. K. Thomas, et al., 2012 Extraordinary genome stability in the ciliate Paramecium tetraurelia. Proc. Natl. Acad. Sci. U. S. A. 109: 19339–19344.

Taylor D. R., C. Zeyl, and E. Cooke, 2002 Conflicting levels of selection in the accumulation of mitochondrial defects in Saccharomyces cerevisiae. Proc. Natl. Acad. Sci. U. S. A. 99: 3690–3694.

Virdi K. S., J. D. Laurie, Y.-Z. Xu, J. Yu, M.-R. Shao, et al., 2015 Arabidopsis MSH1 mutation alters the epigenome and produces heritable changes in plant growth. Nat. Commun. 6: 6386.

Waneka G., A. K. Broz, F. Wold-McGimsey, Y. Zou, Z. Wu, et al., 2024 Disruption of recombination machinery alters the mutational landscape in plant organellar genomes. bioRxivorg.

Wang Y., and D. J. Obbard, 2023 Experimental estimates of germline mutation rate in eukaryotes: a phylogenetic meta-analysis. Evolution Letters 7: 216–226.

Weng M.-L., C. Becker, J. Hildebrandt, M. Neumann, M. T. Rutter, et al., 2019 Fine-grained analysis of spontaneous mutation spectrum and frequency in Arabidopsis thaliana. Genetics 211: 703–714.

Wolfe K. H., W. H. Li, and P. M. Sharp, 1987 Rates of nucleotide substitution vary greatly among plant mitochondrial, chloroplast, and nuclear DNAs. Proceedings of the National Academy of Sciences 84: 9054–9058.

Wu Z., G. Waneka, A. K. Broz, C. R. King, and D. B. Sloan, 2020 MSH1 is required for maintenance of the low mutation rates in plant mitochondrial and plastid genomes. Proceedings of the National Academy of Sciences 117: 16448–16455.

Xu Y. Z., M. P. Arrieta-Montiel, K. S. Virdi, W. B. M. de Paula, J. R. Widhalm, et al., 2011 MutS HOMOLOG1 is a nucleoid protein that alters mitochondrial and plastid properties and plant response to high light. Plant Cell 23: 3428–3441.

Xu S., S. Schaack, A. Seyfert, E. Choi, M. Lynch, et al., 2012 High mutation rates in the mitochondrial genomes of Daphnia pulex. Mol. Biol. Evol. 29: 763–769.

Zhu A., W. Guo, S. Gupta, W. Fan, and J. P. Mower, 2016 Evolutionary dynamics of the plastid inverted repeat: the effects of expansion, contraction, and loss on substitution rates. New Phytol. 209: 1747–1756.

Zou Y., W. Zhu, D. B. Sloan, and Z. Wu, 2022 Long-read sequencing characterizes mitochondrial and plastid genome variants in Arabidopsis msh1 mutants. Plant J. 112: 738–755.

